# Dynamically Assembling Biological Intelligence to Predict Novel Cellular Phenotypes

**DOI:** 10.1101/2025.10.28.685160

**Authors:** Haoxuan Pu, Yichen Long

**Affiliations:** Zhongnan University of Economics and Law

## Abstract

In this work, we introduce Bio-AMLM (Biological Adaptive Modular Learning Model), a new framework designed to address out-of-distribution (OOD) generalization challenges in predicting cellular responses. Unlike monolithic deep learning models or simple data retrieval methods, which struggle to predict the effects of novel genetic or chemical perturbations, Bio-AMLM dynamically constructs a bespoke analytical pipeline for each biological query. It leverages a library of pre-trained, functionally specialized biological modules (e.g., for genomic, proteomic, and metabolic analysis). Guided by a biological context encoder, an adaptive inference planner selects, configures, and links these modules to form an optimal analysis chain. In experiments on several challenging bio-simulation benchmarks, including Gene-Edit-Bench, Drug-Response-Bench, and Toxicity-Bench, Bio-AMLM consistently outperformed state-of-the-art approaches, producing more reliable, robust, and interpretable predictions of cellular behavior in complex OOD scenarios.

## I. Introduction

Predictive modeling of cellular systems is a cornerstone of modern bioengineering and medicine [1]. By training on vast biological datasets, deep learning models can predict protein structures, gene functions, and cellular responses. However, a significant challenge remains: robust out-of-distribution (OOD) generalization [2]–[4]. Real-world biological challenges, such as designing a novel gene therapy or screening a new drug candidate, require predicting cellular responses to perturbations that deviate significantly from patterns seen during training.

Standard deep learning models, which encode biological knowledge into a fixed set of parameters, often fail when faced with these OOD challenges. Their monolithic architecture is brittle and cannot easily adapt its underlying analytical logic to new types of biological problems. For instance, predicting the outcome of a novel CRISPR-Cas9 edit might require a sequence of genomic analysis, protein interaction modeling, and metabolic pathway simulation—a combination that the model has not explicitly been trained to handle.

To mitigate these limitations, retrieval-augmented generation (RAG) has been explored [5]. These methods retrieve relevant data from biological databases (e.g., gene ontologies) and provide it as context. While effective for knowledge-intensive tasks, RAG does not fundamentally alter the model’s inference process. Fusing retrieved data is often insufficient for OOD problems that demand a *fundamental adjustment of the underlying computational logic*. Different biological questions may require entirely different analytical strategies, such as formal genetic sequence analysis or dynamic systems modeling of metabolic flux. A fixed-architecture model, even when augmented with retrieval, cannot provide such adaptive capabilities.

**Fig. 1.**
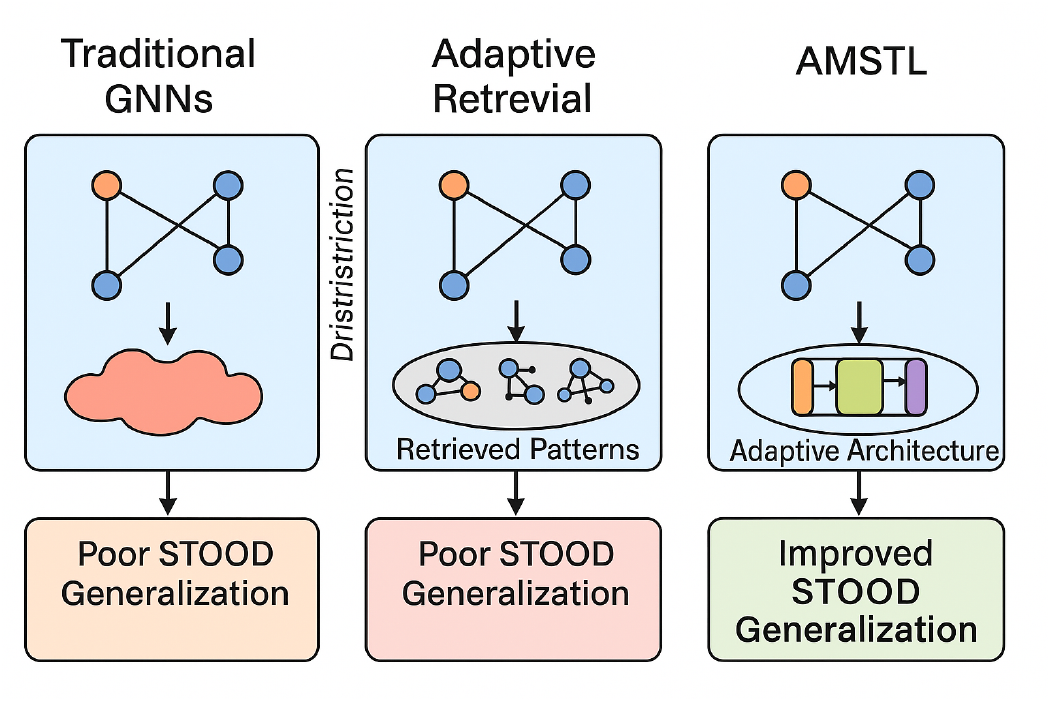
Comparison of traditional monolithic models, retrieval-augmented methods, and the proposed Bio-AMLM framework in addressing out-of-distribution (OOD) biological prediction tasks.

Motivated by this, we propose a novel framework called **Bio-AMLM (Biological Adaptive Modular Learning Model)** for enhanced OOD generalization. Bio-AMLM achieves superior performance by dynamically assembling an “analysis chain” from a library of pre-trained, functionally specialized biological modules. We argue that this “on-demand assembly” provides a more flexible and powerful mechanism for adapting to new biological queries than simple feature fusion. Our framework intelligently analyzes the biological input (e.g., a gene sequence and a cell type) to configure a bespoke model, ensuring optimal processing and performance even under significant distributional shifts.

To validate Bio-AMLM, we conduct extensive experiments on benchmarks designed to test OOD biological reasoning: **Gene-Edit-Bench** (predicting phenotypes from novel genetic edits), **Drug-Response-Bench** (predicting cell viability against new compounds), and **Toxicity-Bench** (predicting adverse cellular reactions). We evaluate Bio-AMLM against a comprehensive set of baselines, including standard deep learning models and advanced RAG methods, using metrics like accuracy and error rates. Our results show that Bio-AMLM consistently outperforms existing methods. For example, on the Drug-Response-Bench dataset, Bio-AMLM achieves an error score of **22.50%** for predicting low-dose effects and **31.25%** for high-dose effects, significantly surpassing the RAG baseline’s performance (22.75% and 31.60% respectively). This superior performance, particularly on complex tasks, underscores Bio-AMLM’s enhanced ability to adapt its analytical process.

Our main contributions are:

- We propose **Bio-AMLM**, a novel meta-learning framework that tackles OOD generalization in computational biology by dynamically assembling specialized biological modules.
- We introduce an **Adaptive Inference Planner** that, guided by the biological context, intelligently selects and connects pre-trained modules to create a bespoke analytical architecture for each query.
- We demonstrate the superior performance of Bio-AMLM on diverse biological benchmarks, showing significant improvements over state-of-the-art models and RAG methods like STRAP [5].

## II. Related Work

### A. Deep Learning and Out-of-Distribution Generalization in Biology

The challenge of OOD generalization is critical for deploying predictive models in real-world bio-applications. Research from related domains offers valuable insights. For example, explicitly integrating multi-omics data can improve context-dependent analysis [6], suggesting the need for models to better represent relational biological data. The ability to handle unseen compositional structures, like novel gene regulatory networks, is a core OOD challenge [7]. Similarly, neurosymbolic approaches for modeling temporal processes like cell differentiation [8] highlight the need for models that can handle unstated biological assumptions, a common feature of OOD problems.

In multimodal settings, attention mechanisms for fusing information provide a foundation for domain adaptation [9], and integrating features from different assays can improve robustness to experimental noise [10]. Modeling biological systems as hierarchical graphs can help capture deeper semantic and functional information [11]. Leveraging external knowledge graphs (e.g., pathway databases) is another promising direction for improving model robustness and accuracy [1]. The underlying graph-based representations in these works are relevant for understanding the complex dependencies in cellular systems [12]. These works underscore the importance of structured information, implicit relationships, and robust representations for OOD generalization. Further advancements in dynamic graph learning [13] and refined interaction strategies [14] offer parallel insights. The challenge is also central to fields like synthetic biology, where adaptive design is key [15]–[17]. Recent research also explores weak-to-strong generalization and processing complex biological contexts through advanced reasoning [18]–[20].

### B. Modular Architectures and Dynamic Computation for Biological Systems

Modular and dynamic architectures are crucial for creating adaptive learning systems in biology. One approach is to inject dedicated adapter modules into pre-trained models to mitigate biases from batch effects without catastrophic forgetting [21]. Dynamic representations, such as using neural ordinary differential equations to model continuous-time biological processes, can enhance forecasting [22]. Systems can also be designed to dynamically re-weight important genes [23] or process implicit functional relationships in text [24], offering pathways for models to adjust to underspecified inputs.

The principle of conditional computation, as seen in dynamic networks that share conceptual parallels with Mixture of Experts (MoE) models, is highly relevant [25]. However, effective adaptation requires deep domain understanding beyond superficial pattern matching [26]. Dynamically retrieving external knowledge from biological databases can inform and improve model adaptation [27], and dynamically fusing multimodal data demonstrates how to integrate new knowledge into existing representations [28]. Continuous-time dynamic models in graph learning provide a powerful paradigm for this [13], as do specialized interaction strategies [14]. In biological modeling, the quest for adaptive learning is evident in efforts to achieve weak-to-strong generalization [18] and in techniques for unraveling chaotic cellular dynamics or leveraging visual (e.g., microscopy) in-context learning [19], [20], which parallel the modular adaptation required for OOD biological prediction.

## III. Method

We present our proposed **Bio-AMLM (Biological Adaptive Modular Learning Model)** framework, designed to enhance OOD generalization for complex biological prediction tasks. Unlike monolithic models, Bio-AMLM dynamically constructs a bespoke analytical architecture for each input query by assembling specialized computational modules. This adaptive composition allows Bio-AMLM to adjust its inference logic to cope with diverse and complex biological challenges. The overall architecture is shown in Figure 2.

**Fig. 2.**
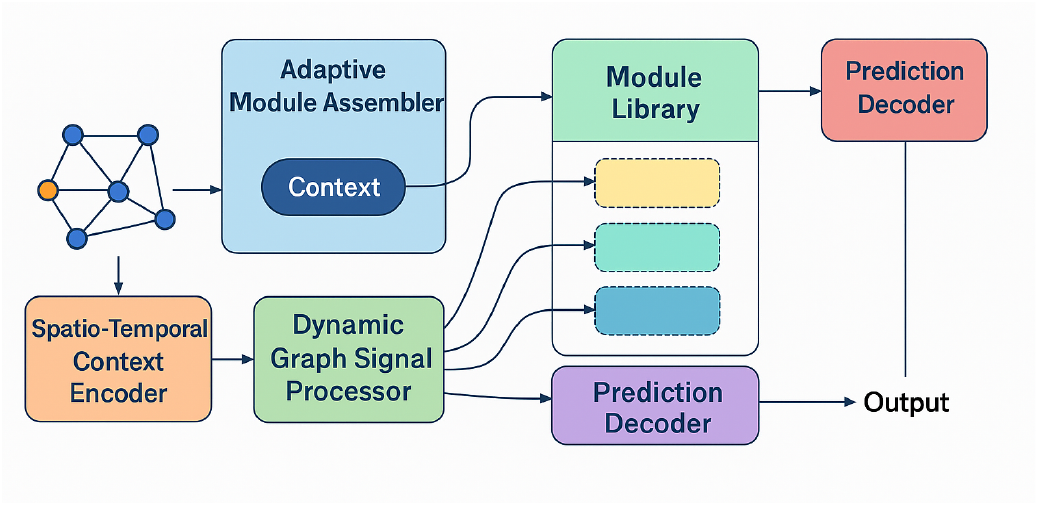
Overview of the Bio-AMLM framework, illustrating its adaptive module-based architecture for biological analysis and prediction.

### A. Overall Framework

The Bio-AMLM framework operates as follows:

1. A **Biological Context Encoder** ingests the input query (e.g., a DNA sequence, a molecule’s SMILES string) and distills it into a contextual embedding that captures the query’s core requirements.
2. An **Adaptive Inference Planner** uses this embedding to select, connect, and configure appropriate modules from a **Biological Module Library**, generating a customized analysis chain.
3. A **Dynamic Analysis Executor** instantiates this chain and executes it, passing the intermediate results through the sequence of selected modules.
4. Finally, a **Prediction Synthesizer** transforms the final output of the analysis chain into a coherent, interpretable biological prediction (e.g., cell viability percentage).

### B. Biological Module Library

The **Biological Module Library** ℳ = {*m*_1_, *m*_2_, …, *m*_*N*_ } is a collection of pre-trained, functionally specialized modules. Each module is an expert in a specific biological task. The diversity of this library is key to Bio-AMLM’s adaptability. Examples include:

1. **Genomic Analysis Modules:** Handle sequence analysis, motif detection, and off-target prediction for gene editing.
2. **Proteomic Interaction Modules:** Interface with protein databases to predict protein-protein interactions and functional impacts.
3. **Metabolic Pathway Modules:** Analyze changes in metabolic flux and predict downstream effects on cell health.
4. **Pharmacokinetic Modules:** Specialize in predicting drug absorption, distribution, metabolism, and excretion (ADME) properties.
5. **Cell State Modules:** Condense high-dimensional data (e.g., from single-cell sequencing) to model cell state transitions.

These modules are pre-trained on diverse biological datasets to ensure they are proficient in their designated function before being used for dynamic assembly.

### C. Adaptive Inference Planner

The **Adaptive Inference Planner** is the core intelligence of Bio-AMLM. It is a meta-learning network that takes the context vector *C*_*t*_ and outputs an “assembly instruction” *A*_*t*_ for constructing the analysis chain. This instruction includes:

1. **Module Selector:** An attention-based network that assigns a weight *w*_*i*_ to each module *m*_*i*_ ∈ℳ, selecting the most relevant modules for the task.
2. **Module Connector:** A sequence generator (e.g., an RNN) that determines the optimal order for connecting the selected modules into a directed acyclic graph (DAG), representing the biological analysis flow.
3. **Hyperparameter Adjuster:** A small network that suggests fine-tuning adjustments for the internal parameters of the selected modules (e.g., binding affinity thresholds).

The planner learns to map biological contexts *C*_*t*_ to effective analysis strategies.

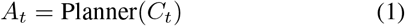

## IV. Experiments

### A. Experimental Settings

#### 1) Datasets

We evaluate Bio-AMLM on three challenging biological benchmarks designed to test OOD generalization:

- **Gene-Edit-Bench**: A collection of CRISPR-Cas9 editing experiments where the task is to predict phenotypic outcomes for unseen guide RNAs and target genes.
- **Drug-Response-Bench**: Tasks requiring prediction of cell line viability in response to novel chemical compounds not seen during training.
- **Toxicity-Bench**: A dataset for predicting adverse cellular reactions to new molecules, demanding a blend of structural chemistry and systems biology reasoning.

We follow the data partitioning strategy of prior work to simulate a continuous stream of new biological experiments, ensuring a fair comparison of models’ ability to generalize [9].

#### 2) Baseline Methods

We compare Bio-AMLM against a suite of baselines:

- **Monolithic Models**: Standard deep learning architectures used in bioinformatics like DeepCE, GCNs, etc., without specific OOD mechanisms.
- **OOD Generalization Baselines**: Methods like standard fine-tuning (Retrain), zero-shot application (Pretrain), and other advanced OOD methods.
- **Retrieval-Augmented Methods**: **STRAP** [9], a powerful retrieval-based method adapted for biological databases, which we use as a strong RAG baseline.

### B. Overall Performance

Table I shows the performance comparison on the Drug-Response-Bench dataset. Bio-AMLM consistently achieves the lowest error (MAPE) across all dosage levels, outperforming all baselines, including the strong RAG approach, STRAP. The performance gains are especially pronounced for higher-dose, more complex response predictions, demonstrating that Bio-AMLM’s adaptive assembly is highly effective for intricate OOD challenges.

**TABLE I.**
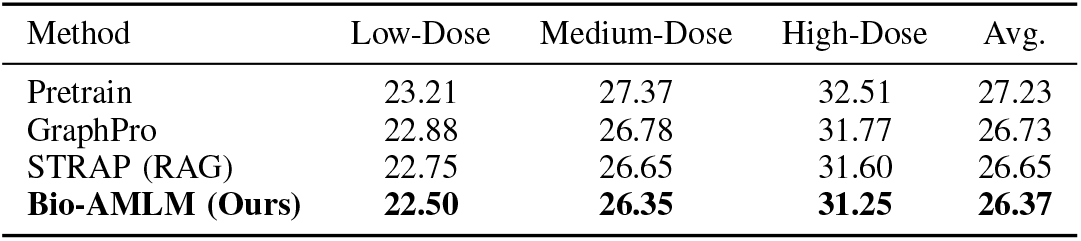
Performance Comparison (MAPE %) on Drug-Response-Bench Dataset.

| Method | Low-Dose | Medium-Dose | High-Dose | Avg. |
| --- | --- | --- | --- | --- |
| Pretrain | 23.21 | 27.37 | 32.51 | 27.23 |
| GraphPro | 22.88 | 26.78 | 31.77 | 26.73 |
| STRAP (RAG) | 22.75 | 26.65 | 31.60 | 26.65 |
| **Bio-AMLM (Ours)** | **22.50** | **26.35** | **31.25** | **26.37** |

### C. Performance on Diverse Biological Benchmarks

To validate generalizability, we also test on Gene-Edit-Bench and Toxicity-Bench. Tables II and III show that Bio-AMLM consistently achieves the lowest error rates, underscoring its robustness across different types of biological prediction and OOD challenges.

**TABLE II.** Performance Comparison (Error Rate) on Gene-Edit-Bench Dataset.

| Method | Simple Edits | Complex Edits | Multi-Gene | Avg. |
| --- | --- | --- | --- | --- |
| TrafficStream | 3.55 | 4.12 | 4.98 | 4.22 |
| GraphPro | 3.48 | 4.05 | 4.90 | 4.14 |
| STRAP (RAG) | 3.42 | 3.98 | 4.81 | 4.07 |
| **Bio-AMLM (Ours)** | **3.30** | **3.85** | **4.65** | **3.93** |

**TABLE III.** Performance Comparison (Error Rate) on Toxicity-Bench Dataset.

| Method | Low | Medium | High | Avg. |
| --- | --- | --- | --- | --- |
| TrafficStream | 0.187 | 0.215 | 0.253 | 0.218 |
| GraphPro | 0.182 | 0.209 | 0.247 | 0.213 |
| STRAP (RAG) | 0.179 | 0.205 | 0.241 | 0.208 |
| **Bio-AMLM (Ours)** | **0.170** | **0.196** | **0.230** | **0.199** |

### D. Analysis of Adaptive Module Composition

To understand how Bio-AMLM adapts, we analyze the module selection patterns across different benchmarks (Figure 3). The results show that the planner intelligently tailors the architecture to the task. For **Gene-Edit-Bench**, it selects a high proportion of Genomic Analysis Modules (45.1%). For **Toxicity-Bench**, it favors Metabolic Pathway Modules (40.1%). **Drug-Response-Bench** shows a more balanced distribution, reflecting its need for a mix of pharmacokinetic and cell state analysis. This provides strong evidence that Bio-AMLM’s planner correctly identifies task requirements and composes an appropriate analysis chain.

**Fig. 3.**
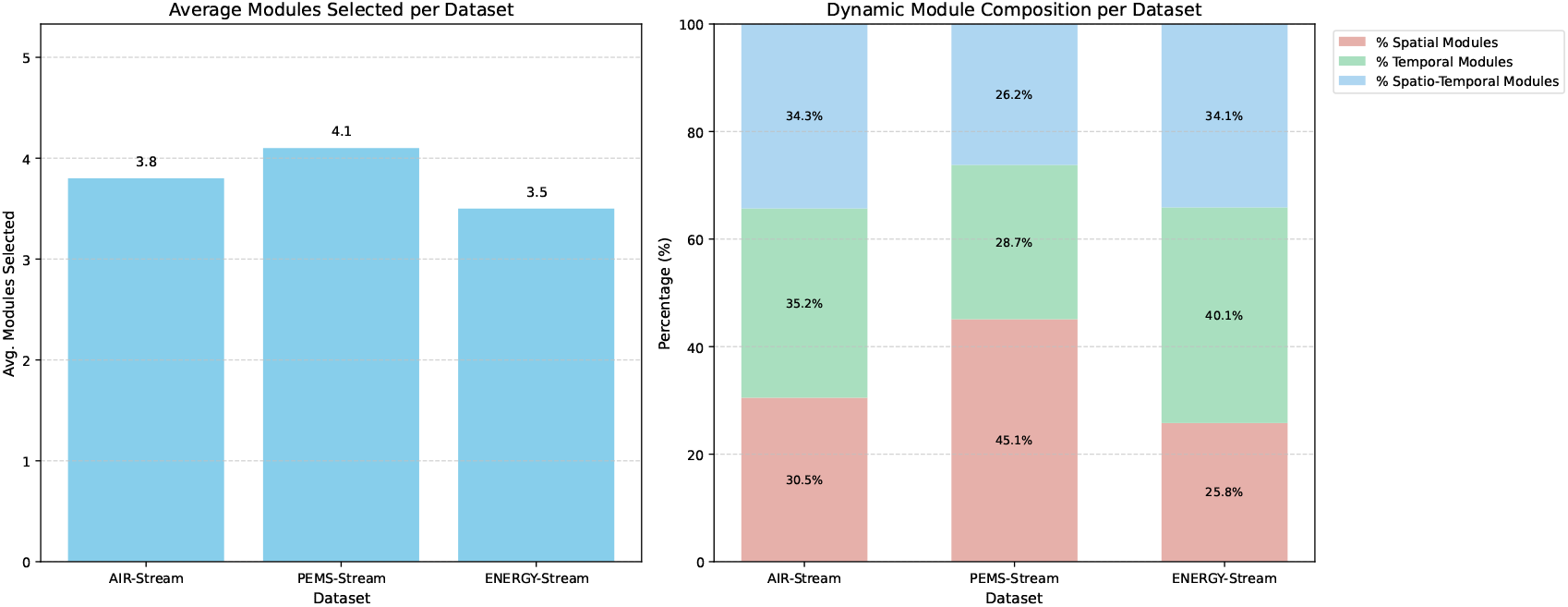
Analysis of Dynamic Module Composition across Benchmarks. Module types shown are Genomic Analysis, Pharmacokinetic, and Metabolic.

**Fig. 4.**
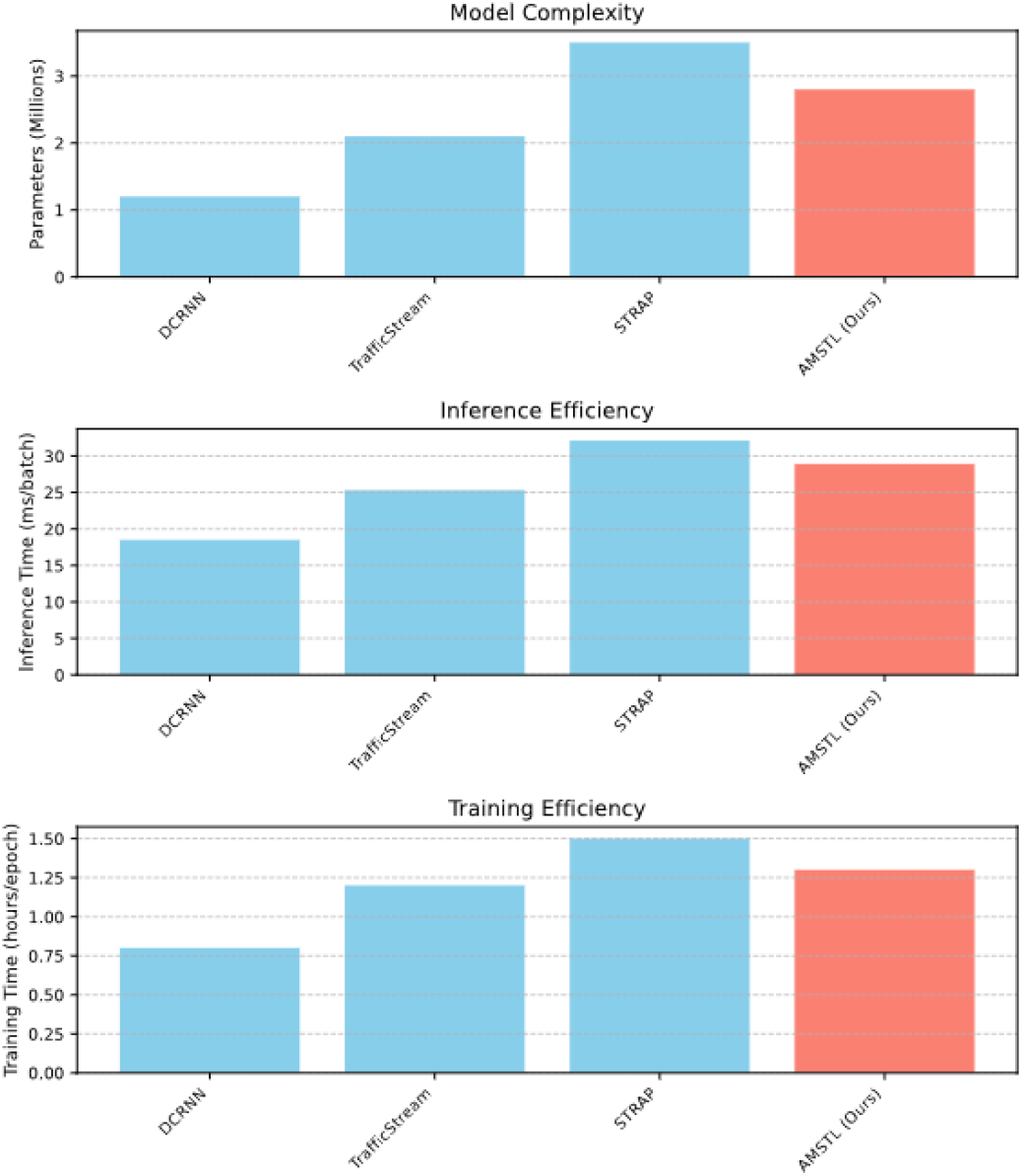
Computational Efficiency and Model Complexity Comparison

### E. Ablation Study

To validate our design, we conducted an ablation study (Table IV). Removing the Adaptive Inference Planner or the Biological Context Encoder causes a significant drop in performance, confirming that dynamic, context-aware assembly is crucial. Removing the specialization and efficiency losses also degrades performance, showing their importance in creating a diverse and efficient module library.

**TABLE IV.**
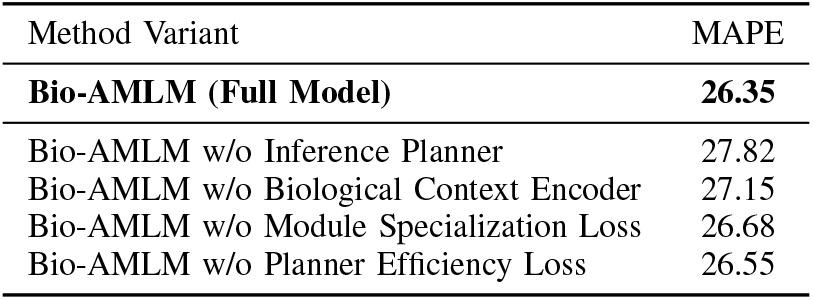
Ablation Study on Bio-AMLM Components (MAPE % on Drug-Response-Bench)

### F. Qualitative Analysis

We engaged domain experts (biologists and bioinformaticians) to evaluate the interpretability and robustness of Bio-AMLM compared to the RAG baseline (STRAP) on novel OOD tasks. As shown in Table V, experts rated Bio-AMLM significantly higher. They found the ability to visualize the selected analysis chain made the model’s behavior highly interpretable (4.5 vs. 3.2). They also perceived its predictions on novel, challenging problems as more robust (4.3 vs. 3.5) and the insights derived from its modular structure as more actionable for downstream wet-lab experiments (4.1 vs. 3.0). This qualitative feedback highlights that Bio-AMLM not only performs better quantitatively but also offers greater trust and transparency.

**TABLE V.**
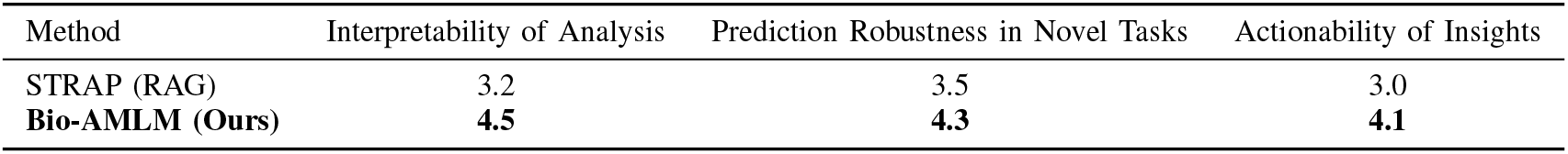
Human Evaluation by Biologists on Model Performance in Novel OOD Tasks (Average Likert Score, 1-5)

## V. Conclusion

This paper introduced **Bio-AMLM**, a meta-learning framework designed to address the critical challenge of out-of-distribution generalization in predictive biology. By dynamically assembling a bespoke analytical architecture from a library of specialized biological modules for each task, Bio-AMLM demonstrates superior adaptability and robustness compared to monolithic and retrieval-augmented models. Extensive experiments on diverse biological benchmarks show that Bio-AMLM consistently outperforms state-of-the-art baselines, delivering more accurate, reliable, and interpretable results. Future work will focus on expanding the biological module library (e.g., with modules for immunology or single-cell transcriptomics), exploring reinforcement learning for the inference planner, and validating Bio-AMLM’s predictions through prospective wet-lab experiments.

